# Polystyrene microplastic exposure disrupts mitochondrial pathways and nuclear processes in primary intestinal epithelial cells

**DOI:** 10.64898/2026.09.22.753448

**Authors:** Charlotte E. Sofield, Anastazja M. Gorecki, Li Shan Chiu, Chidozie C. Anyaegbu, Ryan S. Anderton

## Abstract

Microplastics are pervasive environmental pollutants that are a growing health burden. Humans commonly ingest microplastics orally, yet the resulting proteomic response of the gut epithelium remains unclear. This study investigated the cellular effects of microplastic exposure in primary rat duodenal epithelial cells using proteomics. Cells were exposed to pristine or digested 0.5 µm polystyrene microplastics (MP) at 10 or 100 µg/mL for 72 hours and were then analyzed by tandem liquid chromatography and mass spectrometry (LC-MS). Proteins that were significantly different in intensity compared to controls, with a fold-change of 1.3 or greater, were considered to be differentially expressed. This criterion identified 41 differentially expressed proteins after 100 µg/mL pristine MP exposure, with 10 following 100 µg/mL digested MP exposure. FGSEA pathway analysis revealed that 270 Reactome pathways were significantly altered following microplastic exposure across conditions. These pathways contributed to functional domains including protein synthesis, DNA replication, cell cycle control and aerobic respiration. Overall, microplastic exposure was associated with upregulated mitochondrial respiration, and downregulation of nuclear-related processes including DNA synthesis, transcription and cell proliferation in a dose dependent manner. This study demonstrates the impact of microplastic surface chemistry on biological interactions and provides targets for future investigation (mitochondria and nucleus).

## 1. Introduction

Microplastics are ubiquitous environmental and dietary pollutants, that are of growing concern for human health. Primary microplastics are intentionally produced, largely to be used in cosmetics (e.g. nail enhancement), manufacturing (e.g. 3D printing) and research applications (e.g. flow cytometry calibration). Whereas secondary microplastics are the product of plastic degradation, which primarily occurs due to UV exposure, heat and physical abrasion ^1^. By definition, microplastics are particles smaller than 5 mm in size, although most particles in food and drinking water are smaller than 300 µm^2^. Microplastics spread through food webs, from contaminated water bodies and soil to livestock ingestion and food packaging ^3^, ultimately contaminating the food and water ingested by humans. Consequently, the effects of microplastic exposure in the gastrointestinal system are a critical focus of ongoing research. Clinical studies have identified microplastics in sites throughout the body including the gut ^4^, liver ^5^, kidneys, peripheral vasculature ^6^ and brain ^7^. Emerging pre-clinical evidence indicates that gastrointestinal microplastic exposure is detrimental for cell and tissue health, causing inflammation and histological changes in the intestinal epithelium and mucosa ^8^. These epithelial changes may promote translocation of microplastics from the gut lumen towards peripheral circulation ^9^. Previous studies have shown that microplastic exposure *in vitro* is cytotoxic at high doses and prolonged exposure duration ^10–13^. However, the underlying changes in protein expression driving reported cytotoxicity and cellular effects of microplastic exposure are not well understood.

The intestinal epithelium functions as both an absorptive and protective surface, primarily formed by intestinal epithelial cells. Intestinal epithelial cells comprise the structural back bone of the intestinal barrier, whereby selective permeability across the intestinal epithelium is regulated by intercellular tight junctions ^14^. They are the primary absorptive cell type in the gut lumen and are a highly proliferative population with a relatively short mature life span of 3-7 days. Intestinal epithelial cells are exposed to a number stressors, such as bacterial infection or ischemia, along a spectrum of physiological to pathological conditions ^15^. Under both homeostatic and stressful conditions apoptosis is a critical regulator of intestinal epithelial health and integrity^16^, hence intestinal injury is commonly associated with upregulation of pro-apoptotic proteins^17^ and pro-inflammatory cytokines like interleukin 6 (IL-6) ^18^. Importantly, intestinal epithelial cells have the capacity to recover from mild insults ^18^, and stress does not always result in cell death. Oral microplastic exposure has been consistently associated with an increase in oxidative-stress response proteins and pathways, indicating that microplastics induce survivable stress in intestinal epithelial cells ^19,20^. However, chronic stress in intestinal epithelial cells can lead to barrier dysfunction, inflammation and malignancy ^21^.

We aimed to investigate how microplastic exposure affected the function of primary duodenal epithelial cells using a proteomic approach. To simulate a biologically relevant scenario, we applied an *in vitro* digestion protocol ^22^ to a subset of microplastics prior to exposure in intestinal epithelial cells. Untargeted proteomic analysis revealed that microplastic exposure significantly altered protein expression, disrupting pathways involved in fundamental functions like gene expression and mitochondrial metabolism. Supporting our previous work^22^, this study found that microplastic-induced proteomic changes were partially ameliorated by simulated microplastic digestion prior to cellular exposure.

## 2. Methods

### 2.1 Microplastics

An aqueous suspension of fluorescent 0.56 μm polystyrene microplastics (Purple (555 nm), 0.4-0.6 μm, 1 % w/v in deionized water) was purchased from Spherotech (USA; FP-0562-2, FH-2052-2), as previously described ^22^. Microplastic stock (10 mg/mL) was diluted to 6 mg/mL in sterile PBS prior to use.

### 2.2 Simulated Microplastic Digestion

Immediately prior to experimentation, a modified INFOGEST protocol ^23^ was used to simulate bio-corona formation on microplastics in the digestive tract and was performed as previously described ^22^. Briefly, simulated gastric fluid was combined 1:1 with 6 mg/mL microplastic suspension or PBS and incubated for 2 hours at 37°C. The simulated gastric fluid was then adjusted to pH 7 ± 0.5, added to the simulated intestinal fluid (15 mg/mL bovine serum albumin in PBS to approximate intestinal total protein concentration in a fed state ^24^) at a ratio of 1:2, then incubated for 18 hours at 37°C. This process yielded a suspension of digested microplastics at a concentration of 1000 µg/mL and an aliquot of complete digestive fluid not containing microplastics. To keep media conditions consistent between all samples pristine microplastics were diluted in digestive solution immediately prior to exposure.

### 2.3 Primary intestinal cell culture

Primary duodenal epithelial cells from Sprague-Dawley rats (RN-6047; Cell Biologics) were resurrected into a 25 cm^2^ flask that had been pre-coated with 0.1 % gelatin solution (CB6650; Cell Biologics). Cells were maintained in complete epithelial cell media (M6621; Cell Biologics) containing 10% FBS and 1 % penicillin/streptomycin (GibcoTM; 15070063), in a CO_2_ incubator (5% CO_2_, 65% air balance, 68% humidity, 37°C). Flasks were passaged at 60% confluency using trypsin containing 0.25% EDTA (TrypLE; Gibco) and expanded once prior to plating for experimentation. For experimentation, cells were seeded into 24 well plates pre-coated with poly-D-lysine (10 mins RT, then aspirated) at 8 × 10^5^ cells/ well in complete epithelial cell media with 5 % FBS.

### 2.5 Proteomic analysis of intestinal cells

When epithelial cells were approximately 60 % confluent, culture media was aspirated and replaced with 450 µL of fresh epithelial cell media supplemented with 5% FBS and 50 µL of digestive solution (control), or 50 µL one of four microplastic solutions: pristine and digested, with the final concentration being 10 µg/mL or 100 µg/mL. All cells received the same concentration of complete digestive solution with or without microplastics. 72 hours after exposure, media was aspirated from all wells and cells were rinsed twice with cold PBS on ice for 5 minutes to wash any lightly adhered plastics.

#### 2.5.1 Extraction & sample preparation

Protein lysates were collected and prepared for liquid chromatography and mass spectrometry (LC-MS). Briefly, 100 µL of extraction buffer containing 7% sodium dodecyl sulfate, 0.1 M Tris-HCl pH 7.65, 5 mM TCEP pH 7.5 stabilized Bond-Breaker™ TCEP Solution Neutral pH Pierce (Thermo Scientific) and 1× cOmplete™ protease inhibitor cocktail (Roche), was added to each well. Cells were collected from each well, and the extraction buffer from triplicate wells was pooled in a microcentrifuge tube, on ice. All samples were immediately transferred to -80 °C for storage.

#### 2.5.2. Protein Digest & Cleanup

Protein digest and cleanup was performed by technicians at Centre for Microscopy Characterisation and Analysis (University of Western Australia). Total protein in the cell lysates was quantified using an Amidoblack Protein Quantification Assay, as per the standard protocol ^25^. In preparation for the digest, 60 µg of cell lysate proteins were normalized to equal volumes by the addition of extraction buffer. Proteins were alkylated by adding iodoacetamide to a final concentration of 20 mM, incubating away from light for 30 minutes. Excess iodoacetamide was quenched with 5 mM Dithiothreitol. Proteins were bound to Sera-Mag SpeedBeads (Cytiva) at a 1:8 bead-to-protein ratio in the presence of 50% (v/v) ethanol. After a 10-minute incubation at room temperature with shaking (1000 rpm), samples were placed on a magnetic rack and washed five times with 80% ethanol to remove contaminants. On-bead digestion was performed in 50 mM ammonium bicarbonate at 37 °C, using sequencing-grade trypsin (Promega) at a 1:50 enzyme-to-protein (v/w) ratio for 18 hours. Peptides were recovered from the supernatant after magnetic separation and transferred to LC-MS vials for analysis

#### 2.5.3 LC-MS

LC-MS was performed by Protein digest and cleanup was performed by technicians at CMCA (the WA Proteomics facility). Protein samples were injected into an online nanoflow (0.8 µL / min) capillary column (Picofrit with 50 μm tip opening / 75 μm diameter, New Objective, ICT36015030F-50) packed in-house with 10 cm C18 silica material (3 µm) connected to Thermo Astral, in-line with Dionex Ultimate 3000 series UHPLC. Mobile phase A was 1% formic acid (FA) in water; B was 0.1% FA in acetonitrile (LC-MS grade). The following gradient was used for peptide separation: from 2% B to 6% B over 0.3 min, then to 23% B over 15 min, followed by 35% B over 1 min, and to 80% B over 0.2 min, followed by 0.4 min at 80% B. The column was then equilibrated at 2% A for 2.5 mins prior to the next injection. Eluting peptides were analyzed on a Thermo Orbitrap Astral mass spectrometer operated in positive ion mode using DIA: MS1 scans at 240k resolution (m/z 380–680), AGC target of 500%, and max injection time of 5ms, followed by nonoverlapping DIA windows of 2m/z covering m/z 150–2,000, AGC target of 500% and max precursor accumulation time of 3 ms. MS QC and digest QC samples passed performance thresholds. Raw files were processed with DI-Ann ^26^ and its default parameter. A database consisting of protein sequences from the taxonomy ID 10116 (*Rattus norvegicus*) was used with the following parameters for protein identification: 20 ppm mass tolerance for peptides and 8 ppm mass tolerance for fragments, fixed modifications: carbamidomethyl-cysteine, variable modifications: oxidation of methionine and protein-N-acetylation.

### 2.6 Data Analysis

All samples were plated in triplicate wells, and all experiments were independently repeated three times (n = 3). Raw protein data was imported into in DIA-Analyst (v0.1-.5; Monash University) for MinDet imputation. For pathway analysis, a Reactome pathways data base ^27^ were imported into R version 4.5.1 ^28^ and R studio ^29^. As per the standard method for pre-ranked GSEA analysis, proteins were sorted by strength and significance of change compared to control, using a ranking statistic that was calculated by absolute log_2_ (fold change) multiplied by log_10_ (adjusted p)^30^. Rat genes were then converted to human orthologs for fast gene set enrichment analysis (FGSEA). For FGSEA, upregulated and downregulated pathways were mapped to top parent pathways prior to visualization. To identify the protein drivers of pathway changes, leading edge proteins were extracted from FGSEA output, and were ranked based on influence score which is calculated by:

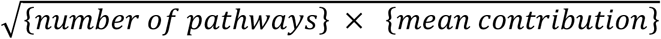

Where mean contribution accounts for fold change and significance:

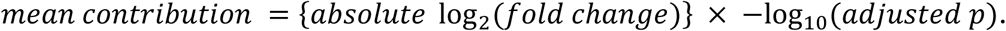

The threshold for differentially expressed proteins and pathway significance was *p* ≤0.05. Leading edge proteins were tested for normality and homogeneity of variety using Levene’s and Shapiro-Wilk tests and then compared by ANOVA and Cohen’s D post hoc test, where the significance threshold was *p* ≤0.05.

## 3. Results

### 3.1 Differentially expressed proteins associated with microplastic exposure

To investigate the intracellular effects of microplastic exposure in the gut epithelium, primary intestinal epithelial cells were exposed to pristine and digested microplastics for 72 hours and then analyzed by untargeted proteomics. Proteomic analyses identified 9148 proteins in total, of which 132 unique proteins were significantly altered, with 15 significant protein changes in cells exposed to 100 µg/mL digested MP samples, and 131 significant protein changes in cells exposed to 100 µg/mL pristine MP. No duplicated protein identities were present, but 13 unvalidated proteins were removed ^31^.

For exploratory analysis, a fold change threshold of 1.3 was set for significantly modified proteins, which yielded 41 differentially expressed proteins (refer to supplementary material for a complete list of proteins). Differentially expressed proteins were visualized across exposure conditions using a heat map, illustrating that microplastic exposure induced bi-directional changes in protein expression (Figure 1). In cells exposed to 100 µg/mL pristine MP, 19 differentially expressed proteins were downregulated and 21 were upregulated, whereas following exposure to 100 µg/mL digested MP, 3 differentially expressed proteins were upregulated and 7 were down regulated. Nine differentially expressed proteins were shared between digested and pristine 100 µg/ml exposure conditions. Of these, solute carrier family 25 member 33 (slc25a33) and pigment epithelium-derived factor (Serpinf1) were upregulated. Shared downregulated proteins included cytochrome c oxidase subunit (Ndufa4), oxidized low-density lipoprotein receptor 1 (Olr1), catalase (Cat), cytochrome c oxidase subunit 4 (Cox4i1), methionine aminopeptidase 2 (Metap2), procollagen, type V, alpha 2 (col5a2) and peroxisomal phytanoyl-CoA dioxygenase (Phyh). In contrast, no differentially expressed proteins were identified after exposures at lower MP doses.

**Figure 1.**
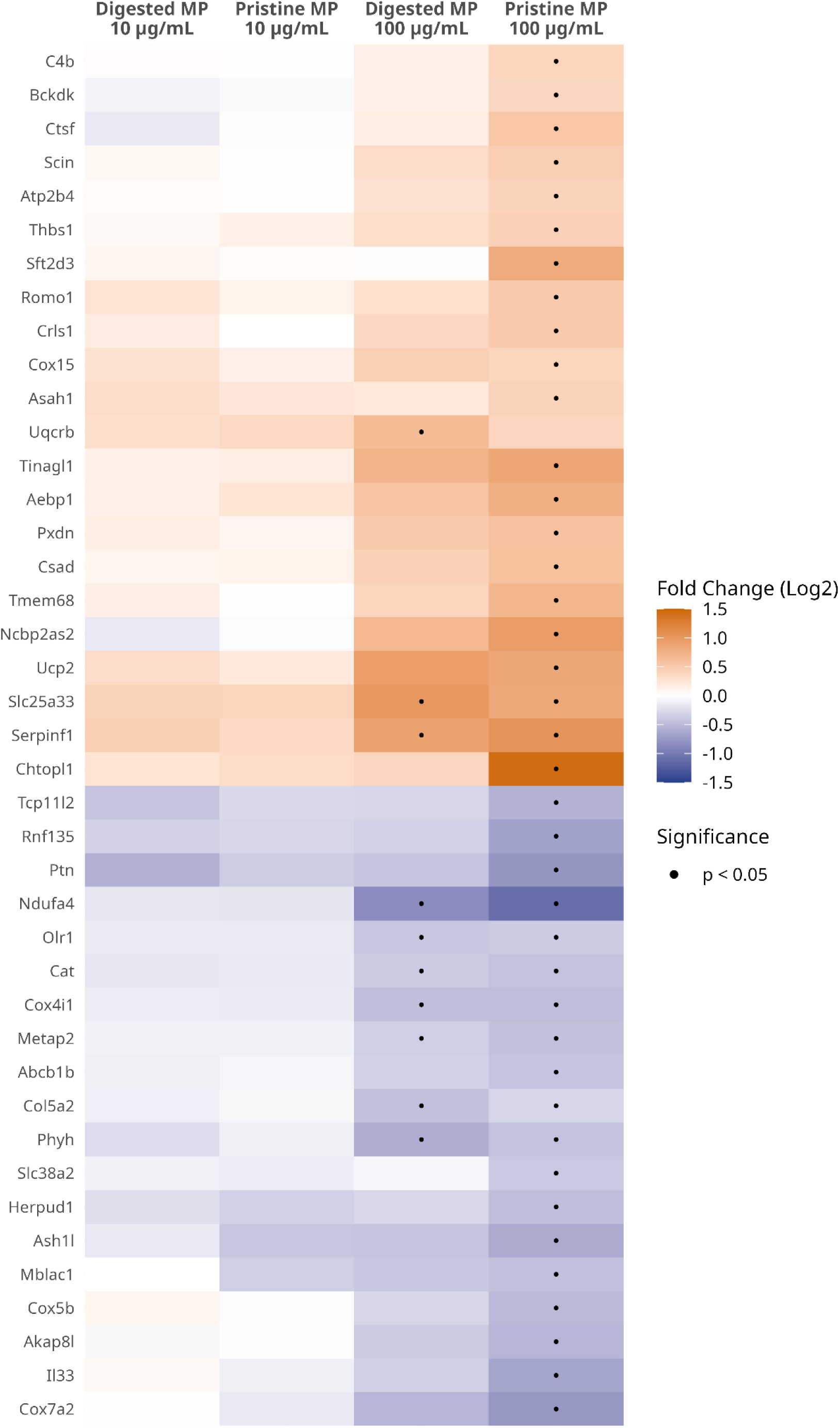
Differentially expressed proteins in primary intestinal epithelial cells exposed to microplastics. Heat map represents significantly altered proteins (n =40; *p* < 0.05) with a fold change of > 1.3, clustered by Euclidean distance and Ward’s method for visualization. No differentially expressed proteins were identified at the 10 µg/mL dose. At the higher dose, pristine microplastics were associated with a greater number of differentially expressed proteins than digested microplastics. Refer to supplementary material for full protein names.

### 3.2 Fast gene set enrichment analysis reveals complex pathway changes associated with microplastic exposure

FGSEA is a pathway analysis tool that considers the entire protein set when describing the cumulative effect of many small or non-significant protein changes on cellular function^32^. Importantly, FGSEA can describe both the direction and magnitude of pathway enrichment. FGSEA yielded 270 unique significantly altered pathways (*p* < 0.049); 221 for 100 µg/mL pristine MPs, 187 for 100 µg/mL digested MPs, 30 for 10 µg/mL pristine MPs and 44 for 10 µg/mL digested MPs, when compared to control cells. Notably, FGSEA identified significant pathway alterations at 10 µg/mL despite an absence of differentially expressed proteins at this concentration. Significantly altered pathways were sorted according to absolute normalized enrichment score, and the top 50 pathways were visualized (Figure 2). FGSEA indicated that microplastic exposure caused significant moderate-strong upregulation (NES = 1.83 – 2.28) or downregulation (NES = - 1.46-(−2.62)) of pathways involved in metabolism of RNA, metabolism of proteins, developmental biology, cell cycle, metabolism, gene expression, disease, cellular responses to stimuli, protein localization and vesicle mediated transport compared to control cells. Overall, the pathway analyses revealed that cellular processes associated with gene expression and protein synthesis were broadly affected by microplastic exposure in a dose dependent manner. Interestingly, pathways appeared to be similarly affected between pristine and digested microplastics in the high dose.

**Figure 2.**
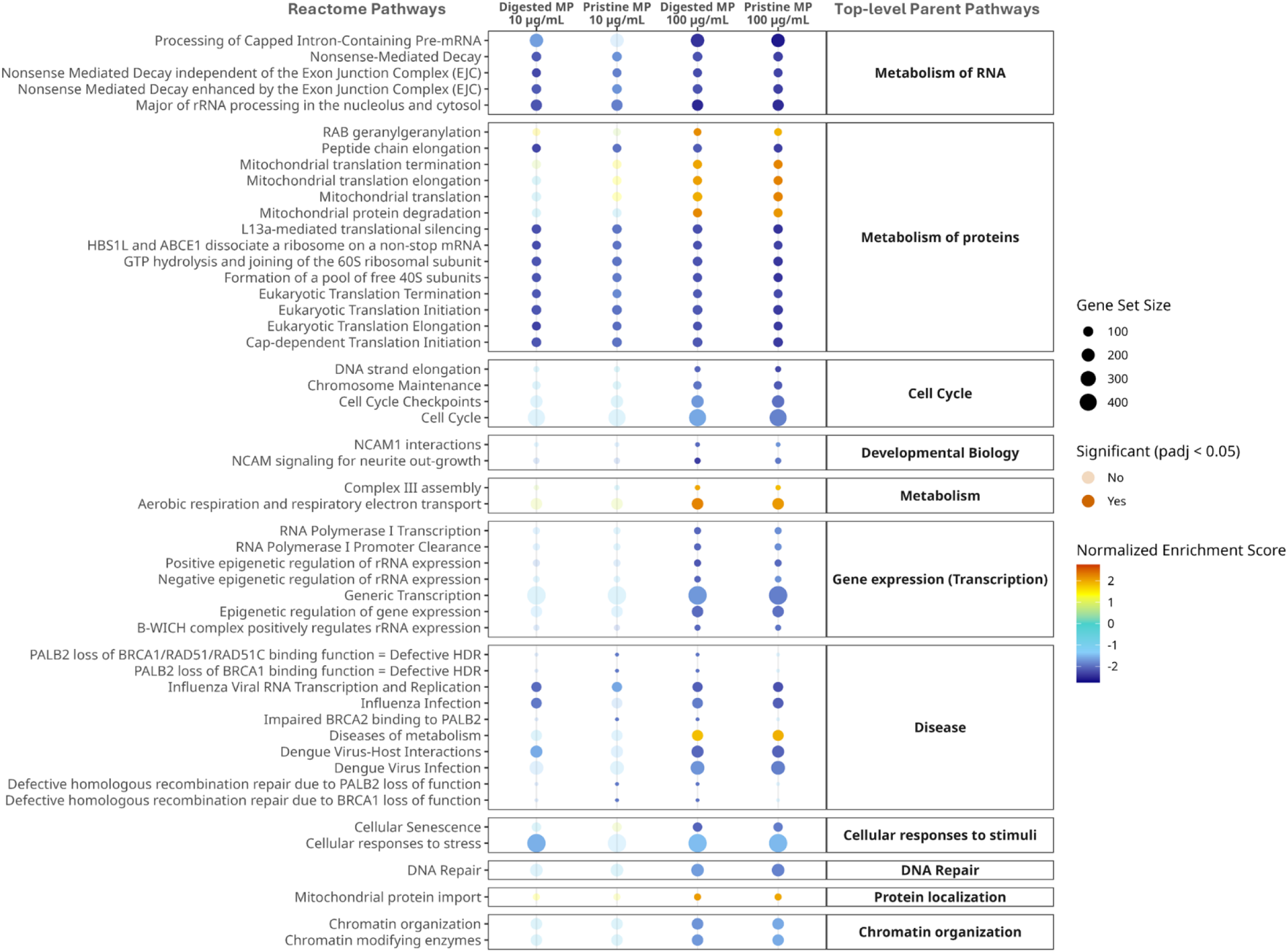
Microplastic exposure resulted in differential pathway enrichment and depletion in primary intestinal epithelial cells. Top 50 significantly (p < 0.05) altered Reactome sub-pathways following microplastic exposure according to FGSEA. Sub-pathways (left y-axis) were grouped in facets according to their top-level parent pathways. Top-level pathway facets are arranged in order of maximum absolute normalized enrichment score (NES) magnitude. As per the legend; color correlates to NES (orange = upregulated, turquoise = no change, blue = downregulated, dot size indicates the number of genes contributing to each pathway, while opacity represents pathway enrichment significance for each exposure condition.

### 3.3 Leading edge proteins associated with gene transcription drive microplastic induced pathway changes

To probe the source of microplastic-induced pathway changes, leading edge proteins were extracted from significant pathways identified by FGSEA. Leading edge proteins form the core protein set that accounts for normalized enrichment scores of upregulated and downregulated pathways. Simply, these are the proteins that drive strong, significant pathway changes. In this data set, leading edge proteins were ranked based on influence score, meaning that the top ranked leading-edge proteins exhibited the greatest combination of fold change, significance and breadth of pathway involvement. To visualize the functional core of microplastic associated effects, leading edge proteins were mapped against their pathway involvement (Figure 3). Following 100 µg/mL pristine MP exposure, the highest influence score was held by replication protein A 32kDa subunit (RPA2; 7.13), closely followed by DNA polymerase delta subunit 2 (POLD2; 6.91), proliferating cell nuclear antigen (PCNA; 6.47), DNA-directed RNA polymerase subunit (POLR2G; 5.58) and cyclin-H (CCNH; 5.42) (Figure 3A). The top 5 leading proteins for 100 µg/mL digested MP were CDK-activating kinase assembly factor MAT1 (MNAT; 5.09), serine-protein kinase ATM (ATM; 4.83), small ribosomal subunit protein eS10 (RPS10; 4.82), replication factor C subunit 4 (RFC4; 4.59) and small ribosomal subunit protein uS10 (RPS20; 4.58) (Figure 3B). This mapping indicated that leading edge genes were largely involved in processes associated with DNA synthesis and repair, DNA transcription, cell cycle regulation and protein synthesis. The intensity of these leading-edge proteins was compared between exposed and unexposed cells, showing that all were significantly expressed in at least one experimental group (Figure 3C). Importantly, significantly identified proteins demonstrated large effect sizes (1.865 – 6.319) (Table S2), indicating that despite exhibiting a small fold change, these proteins were important drivers of microplastic-associated functional changes in intestinal epithelial cells.

**Figure 3.**
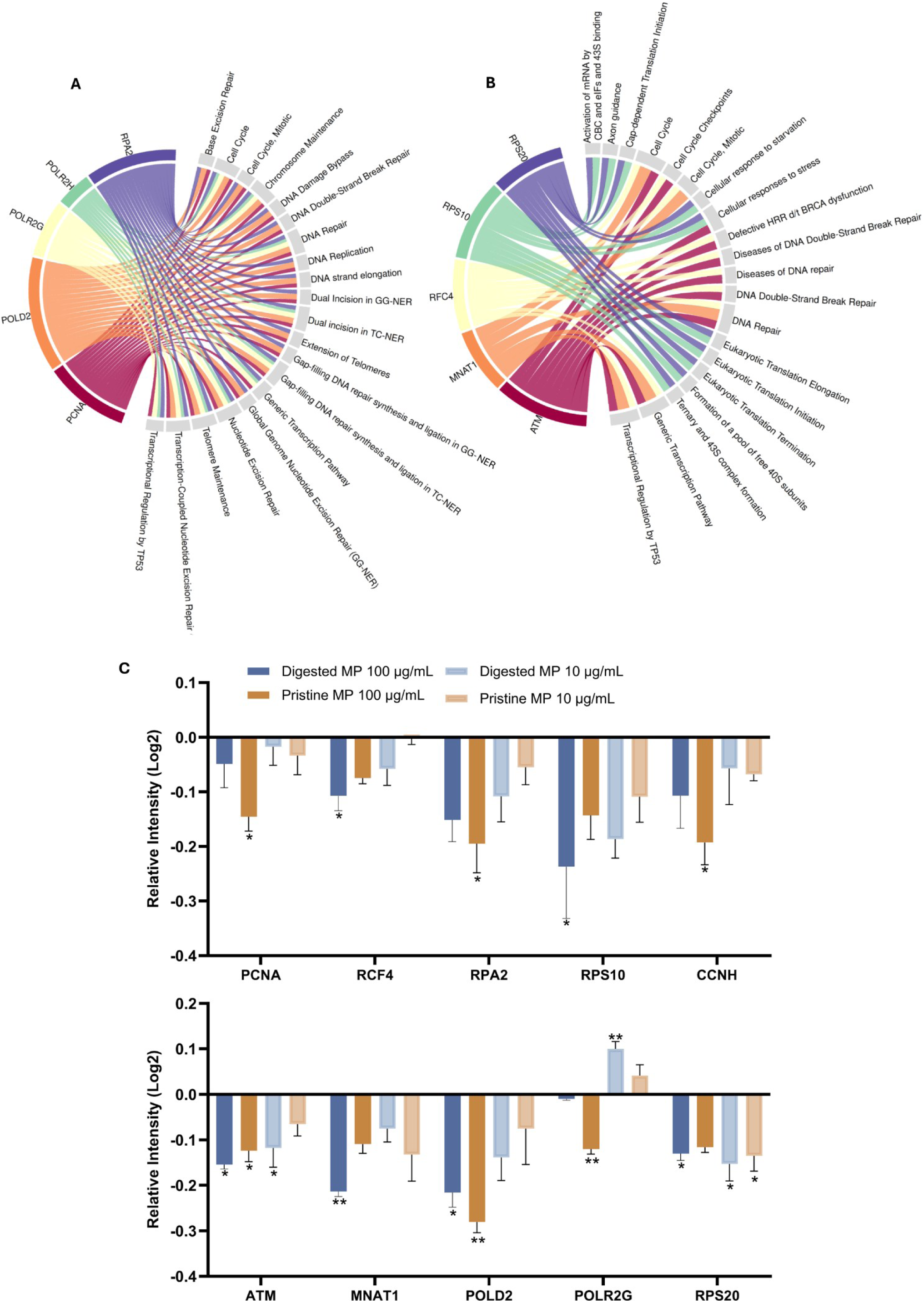
Proteins involved in DNA replication drive microplastic-induced pathway changes. Chord plot of **A)** pristine 100 µg/mL MP and **B)** digested 100 µg/mL MP data connects leading edge proteins, as ranked by influence score (top 20 pathway hits). Refer to supplementary material for full protein names. For both conditions the primary functions of leading-edge proteins were centered around DNA repair and DNA replication, cell cycle control and protein synthesis. **C)** Relative signal intensity of significant leading-edge proteins (p < 0.05), PCNA, RCF4, RPA2, RPS10, CCNH, ATM, MNAT1, POLD2, POLR2G and RPS20 compared to unexposed controls. Significance tested by ANOVA and Dunnett’s post hoc test.

## 4. Discussion

Proteomic analyses revealed that exposure to pristine and digested 100 µg/mL MP resulted in bidirectional differential protein expression. Our selected microplastic concentrations were informed by previous studies ^13,33,34^, designed to exert effects within the time constraints associated with culturing a proliferative cell line. Although the upper microplastic dose is higher than estimated background exposure for humans (e.g. 0.00002–656.8 μg/L in drinking water ^2^), certain ‘high risk’ exposure events (e.g. reheating food in a plastic takeaway container ^35,36^ or tea bag use) may result in transient peaks in microplastic exposure. The accumulation of these high risk events over the course of a day, or several days (given that gut transit time is commonly 30-40 hours), will likely contribute to cumulative luminal microplastic exposure that is far greater than background estimates ^4^. Post Benjamini-Hochberg adjustment, no significant protein changes were associated with exposure to 10 µg/mL microplastics in this model and analysis. This dose dependent effect aligned with prior studies which observed significant microplastic toxicity at higher dose ranges (50 -200 µg/mL), and after 72 hours of exposure ^33,34^.

To mimic elements of the luminal environment, a subset of microplastics were artificially digested prior to exposure to intestinal epithelial cells. Interestingly, microplastic digestion resulted in a lower number of differentially expressed proteins, compared to pristine microplastics in the upper dose. This aligned with our recently published work ^22^, which showed that simulated microplastic digestion significantly reduced microplastic-cell interactions. Despite this phenomenon, differentially expressed proteins common to both groups were identified. These proteins were involved in mitochondrial respiration and cell proliferation. Considering these findings, we used pathway analysis to explore the functional implications of microplastic-associated variations in protein expression. Pathway analysis revealed significantly altered pathways across multiple functional domains. FGSEA considers the whole protein landscape for pathway analysis, meaning that significant pathway alterations could be identified for all exposure conditions. However, fewer significant pathways were identified for the 10 µg/mL conditions than the 100 µg/mL conditions. Pathways of relevance and common across doses were those involved in mitochondrial metabolism, DNA organization, replication and repair and protein synthesis. Given that we studied intestinal epithelial cells, pathways falling under developmental biology and disease were beyond the scope of this study. However, it is noteworthy that neural cell adhesion molecule (NCAM) variants (e.g. L1CAM) are expressed in the intestine, are required for normal enteric development and are implicated in colonic cancer ^37^. However, these proteins were not significantly altered in our data set. Taken together, our findings indicate that microplastic exposure had two main sites of impact in primary intestinal epithelial cells: mitochondria and the nucleus.

Differentially expressed protein analysis indicated that microplastic exposure disrupted homeostatic mitochondrial function, particularly affecting aerobic respiration. Previous work has linked microplastic exposure with abnormal mitochondrial morphology, mitochondrial membrane impairment and mitophagy ^38–40^. Interestingly, we found that peroxisomal phytanoyl-CoA dioxygenase (PHYH) was downregulated by microplastic exposure, indicating that α-oxidation of fatty acids was inhibited. Prior research indicates that microplastics disrupt mitochondrial respiration through upstream suppression of glycolysis ^40^, altered expression of electron transport chain (ETC) complex proteins ^39^ and reduced oxidative phosphorylation. Reflecting this disruption, we observed upregulation of mitochondrial membrane protein ubiquinol-cytochrome c reductase binding protein (UQCRB), which is involved in ubiquitin binding and electron transfer ^41^, and solute carrier family 25 member 33 protein (slc25a33), which is involved in oxidative phosphorylation. Conversely, we observed downregulation of two cytochrome C oxidase (complex IV) subunits (Ndufa4, Cox4i1), which are required for ETC catalysis ^42,43^. Pathway analysis revealed that several mitochondrial protein synthesis processes were upregulated in response to microplastic exposure, as was mitochondrial protein import. Accordingly, aerobic respiration and ETC were upregulated—this finding agrees with a previous study of prolonged microplastic exposure in epithelial cells ^44^ and possibly reflects an increase in O_2_ consumption, which is part of the mitochondrial endoplasmic reticulum stress response used to mitigate ROS induced damage ^45^. Although another study using similar concentrations to our work found that 48 hours of neuronal nanoplastic exposure depressed aerobic respiration overall, our results suggest that microplastic-associated disturbance of mitochondrial respiration is dependent on cell type and the specific conditions of exposure. Disturbed mitochondrial respiration may lead to depletion of intracellular ATP, ETC electron leakage and increased ROS production ^38–40^. Consequently, microplastic exposure has been consistently associated with protein mis-folding ^20^ and upregulation of ROS-mediating enzymes ^19^, such as pigment epithelium-derived factor (Serpinf1) which reduces oxidative stress through mitochondrial stabilization and ROS-associated enzymes, and was found to be upregulated by microplastic exposure ^46^. Unexpectedly, known ROS-mediator proteins catalase (CAT) and oxidized low-density lipoprotein receptor 1 (Olr1) ^47^ were downregulated following microplastic exposure.

Differentially expressed proteins and pathway analysis indicated that DNA synthesis, transcription and cell proliferation were other important functions affected by microplastic exposure in intestinal epithelial cells. For example, upregulated pigment epithelium-derived factor suppresses cell proliferation. Conversely, pro-proliferative proteins ^48,49^ methionine aminopeptidase 2 (Metap2) and procollagen, type V, alpha 2 (col5a2) were both downregulated by microplastic exposure. The upregulation of anti-proliferative proteins and downregulation of pro-proliferative proteins suggests that cell division and replacement were reduced following microplastic exposure. This was corroborated by pathway and leading-edge protein analysis which revealed that transcription, translation, and DNA damage responses were core functions of significantly downregulated leading-edge proteins in the high dose microplastic exposure conditions. For example, replication protein A (RPA2) ^50^, polymerase II subunit G (POLR2G) ^51^, proliferating cell nuclear antigen (PCNA) ^52^, replication factor C subunit 4 (RFC4) ^53^ and DNA polymerase delta 2 (POLD2) ^54^ were significantly down regulated by high microplastic exposure. These proteins are critical for homeostatic DNA replication and transcription and are necessary for DNA damage repair responses. Similarly, serine protein kinase (ATM) ^55^, Cyclin H (CCNH) and CDK-activating kinase assembly factor MAT1 (MNAT1) ^56^ were significantly down regulated. ATM senses DNA damage and ROS, activates double strand repair responses ^55^ and activates checkpoint signaling, while MNAT1 and CCHN form the cyclin-dependent kinase-activating complex which regulates cell cycle progress via downstream RNA polymerase II activation respectively ^56^. Additionally, small ribosomal subunit protein uS10 (RPS20) and 40S ribosomal protein 10 (RS10) were both significantly downregulated, reflecting the observed downregulation of eukaryotic translational pathways ^31^.

Overall, our results suggest microplastics inhibit cellular mechanisms of intestinal epithelial cell proliferation, a particular concern given the role of intestinal epithelial cells in nutrient absorption and barrier function. The cellular dysfunction identified herein may underly microplastic-associated impairment of gut barrier integrity, which has been evidenced in mice ^57,58^ and has systemic consequences ^59^. Notably, similar inhibition of DNA replication and cell proliferation in intestinal epithelial cells has been observed in response to other pollutants (polychlorinated biphenyl 153) ^60^, food additives (e.g. aspartame) ^61^ and ageing ^62^. While our primary culture model did not demonstrate strong evidence of inflammatory activation in response to microplastic exposure, future research including immune-active cells is warranted. Overall, our findings indicate that microplastics may affect proliferative cells by fundamentally impairing their ability to replicate. However, cell proliferation was not specifically quantified in this study, and should be investigated further.

The present study measured cellular outcomes of acute microplastic exposure in the intestinal epithelium; however, it should be noted that humans are exposed chronically to microplastics. The intestinal epithelium exhibits resilience against mild to moderate insults such as pathogen associated molecular patterns (e.g. lipopolysaccharide) and nutrient depletion ^18^. However, over time, the accumulation of endogenous and environmental stressors can lead to clinically significant intestinal dysfunction. For example, prolonged intestinal inflammation is associated with emergence of motor symptoms in models of Parkinson’s disease ^63^, while chronic psychological stress is thought to increase the risk of flare-ups in individuals with inflammatory bowel disease ^64^. Our findings indicate that even acute microplastic exposure in an *in vitro* setting disrupts intestinal epithelial cell homeostasis, which likely contributes to broader intestinal dysfunction. Importantly, microplastic exposure is one of many insults that the intestinal barrier is faced with as part of routine daily exposure. Therefore, we suggest that microplastic exposure might cause primary cellular disruption and may also compound the influence of other endogenous and exogenous disruptors in the gut. For example, microplastics have been shown to promote *Helicobacter pylori* colonization ^65^, increase antimicrobial resistance ^66^, and worsen intestinal injury associated with heavy metal exposure ^67^. Thus, future research should consider the chronic, systemic effects of microplastic exposure in the context of other intestinal challenges.

## Conclusion

This study used a proteomic approach to investigate the functional effects of microplastic exposure in primary intestinal epithelial cells. Our findings indicate that mitochondrial metabolism, DNA replication and transcription, are disturbed by microplastic exposure. We also identified differences in responses to pristine versus digested microplastics, reinforcing the importance in improving the biological relevance of *in vitro* models of microplastic exposure wherever possible. The current work has limitations, such as the lack of detectable responses at lower microplastic doses. Subtle changes occurring at lower concentrations of microplastic exposure may be better captured by a targeted acquisition-analysis pipeline, focusing on protein classes involved in mitochondrial function, DNA replication and cell cycle control. Overall, this study provides candidate pathways for further investigation regarding the acute and chronic implications of microplastic exposure in the intestinal epithelium.

## Acknowledgments

Analysis for this work was performed by CMCA (the WA Proteomics facility) and was supported by infrastructure funding from the Western Australian State Government in partnership with the Australian Federal Government, through the National Collaborative Research Infrastructure Strategy (NCRIS).

## CRediT Author Contribution Statement

Conceptualization; CES, RSA, AMG. Methodology; CES, RSA. Formal analysis; CES. Investigation; CES. Writing-Original Draft; CES. Writing – Review & Editing; CES, CCA, RSA, LSC, AMG. Project administration; CES, RSA, AMG. Resources; CCA, AMG. Supervision; CCA, RSA, LSC, AMG. Funding Acquisition; RSA, AMG. All authors have read and agreed to the published version of the manuscript.

## Data Availability

The data produced by this study is not publicly available due to the size of the data files. Data is available on request to the corresponding author.

