## Supplementary Material for "Polystyrene microplastic exposure disrupts mitochondrial pathways and nuclear processes in primary intestinal epithelial cells"

| **Gene** | **Protein** |
| --- | --- |
| Abcb1b | P-type phospholipid transporter |
| Aebp1 | Adipocyte enhancer-binding protein 1 |
| Akap8l | A kinase (PRKA) anchor protein 8-like |
| Asah1 | Ceramidase |
| Ash1l | Histone-lysine N-methyltransferase ASH1L |
| Atm | Serine-protein kinase ATM |
| Atp2b4 | Calcium-transporting ATPase |
| Bckdk | Protein-serine/threonine kinase |
| C4b | Complement component 4, gene 2 |
| Cat | Catalase |
| Ccnh | Cyclin-H |
| Col5a2 | Procollagen, type V, alpha 2 |
| Cox4i1 | Cytochrome c oxidase subunit 4 |
| Cox5b | Cytochrome c oxidase subunit 5B |
| Cox7a2 | Cytochrome c oxidase subunit 7A2 |
| Cox15 | COX15 homolog, cytochrome c oxidase assembly protein (Yeast) |
| Crls1 | Cardiolipin synthase (CMP-forming) |
| Csad | Cysteine sulfinic acid decarboxylase |
| Ctsf | Cathepsin F |
| Herpud1 | Homocysteine-inducible, endoplasmic reticulum stress-inducible, ubiquitin-like domain member 1 |
| Il33 | Interleukin-33 |
| Mblac1 | Metallo-beta-lactamase domain-containing protein 1 |
| Metap2 | Methionine aminopeptidase 2 |
| Mnat1 | CDK-activating kinase assembly factor MAT1 |
| Ncbp2as2 | NCBP2 antisense 2 |
| Ndufa4 | Cytochrome c oxidase subunit NDUFA4 |
| Olr1 | Oxidized low-density lipoprotein receptor 1 |
| Pcna | Proliferating cell nuclear antigen |
| Phyh | Phytanoyl-CoA dioxygenase, peroxisomal |
| Pold2 | DNA polymerase delta subunit 2 |
| Polr2g | DNA-directed RNA polymerase subunit |
| Ptn | Pleiotrophin |
| Pxdn | Peroxidasin |
| Rfc4 | Replication factor C subunit 4 |
| Rnf135 | E3 ubiquitin-protein ligase RNF135 |
| Romo1 | Reactive oxygen species modulator 1 |
| Rpa2 | Replication protein A 32 kDa subunit |
| Rps10 | Small ribosomal subunit protein eS10 |
| Rps20 | Small ribosomal subunit protein uS10 |
| Scin | Scinderin |
| Serpinf1 | Alpha-2 antiplasmin |
| Sft2d3 | Vesicle transport protein |
| Slc25a33 | Solute carrier family 25 member 33 |
| Slc38a2 | Sodium-coupled neutral amino acid symporter 2 |
| Ucp2 | Uncoupling protein 2 (Mitochondrial, proton carrier) |
| Uqcrb | Cytochrome b-c1 complex subunit 7 |
| Tcp11l2 | T-complex 11 (Mouse) like 2 |
| Thbs1 | Thrombospondin-1 |
| Tinagl1 | Tubulointerstitial nephritis antigen-like |
| Tmem68 | Transmembrane protein 68 |

**Table S1. Protein Naming**

**Table S2. Leading edge protein intensity statistics (relative to control)**

| **Protein** | **Digested MP 100 µg/mL** | **Pristine MP 100 µg/mL** | **Digested MP 10 µg/mL** | **Pristine MP 10 µg/mL** |
| --- | --- | --- | --- | --- |
| ATM | *Effect Size =* -3.962  ***p =* 0.0109*** | *Effect Size =* -2.628  ***p =* 0.036*** | *Effect Size =* -1.865  ***p =* 0.0461*** | *Effect Size =* -1.333  *p =* 0.3509 |
| CCNH | *Effect Size =* -1.458  *p =* 0.3120 | *Effect Size =* -3.771  ***p =* 0.0369*** | *Effect Size =* -0.699  *p =* 0.7725 | *Effect Size =* -3.760  *p =* 0.6660 |
| MNAT1 | *Effect Size =* -6.319  ***p =* 0.0035*** | *Effect Size =* -2.756  *p =* 0.1205 | *Effect Size =* -1.598  *p =* 0.3532 | *Effect Size =* -1.703  *p =* 0.0540 |
| PCNA | *Effect Size =* -0.833  *p =* 0.6818 | *Effect Size =* -3.655  ***p =* 0.0322*** | *Effect Size =* -0.371  *p =* 0.9849 | *Effect Size =* -0.689  *p =* 0.8739 |
| POLD2 | *Effect Size =* -4.031  ***p =* 0.0295*** | *Effect Size =* -6.024  ***p =* 0.0063*** | *Effect Size =* -1.939  *p =* 0.1848 | *Effect Size =* -0.729  *p =* 0.6404 |
| POLR2G | *Effect Size =* -0.404  *p =* 0.9775 | *Effect Size =* -4.257  ***p =* 0.0014*** | *Effect Size =* 3.176  ***p =* 0.0054*** | *Effect Size =* 1.065  *p =* 0.2996 |
| RFC4 | *Effect Size =* -2.225  ***p =* 0.0277*** | *Effect Size =* -1.986  *p =* 0.1381 | *Effect Size =* -1.130  *p =* 0.2958 | *Effect Size =* -0.026  *p =* 1.0000 |
| RPA2 | *Effect Size =* -2.250  *p =* 0.0924 | *Effect Size =* -2.440  ***p =* 0.0283*** | *Effect Size =* -1.463  *p =* 0.2775 | *Effect Size =* -0.909  *p =* 0.7724 |
| RPS10 | *Effect Size =* -2.006  ***p =* 0.0355*** | *Effect Size =* -2.474  *p =* 0.2484 | *Effect Size =* -3.925  *p =* 0.1034 | *Effect Size =* -1.790  *p =* 0.4554 |
| RPS20 | *Effect Size =* -2.603  ***p =* 0.0338*** | *Effect Size =* -2.380  *p =* 0.0589 | *Effect Size =* -2.345  ***p =* 0.0138*** | *Effect Size =* -2.177  ***p =* 0.0277*** |

*Effect size = Cohen’s D ; * Significance = ANOVA + Dunnet Post-hoc*
